# The antimalarial natural product salinipostin A identifies essential α/β serine hydrolases involved in lipid metabolism in *P. falciparum* parasites

**DOI:** 10.1101/827287

**Authors:** Euna Yoo, Christopher J. Schulze, Barbara H. Stokes, Ouma Onguka, Tomas Yeo, Sachel Mok, Nina F. Gnädig, Yani Zhou, Kenji Kurita, Ian T. Foe, Stephanie M. Terrell, Michael J. Boucher, Piotr Cieplak, Roger G. Linington, Jonathan Z. Long, Anne-Catrin Uhlemann, Eranthie Weerapana, David A. Fidock, Matthew Bogyo

## Abstract

Salinipostin A (Sal A) is a potent antimalarial marine natural product with an undefined mechanism of action. Using a Sal A-derived activity-based probe, we identify its targets in the *Plasmodium falciparum* parasite. All of the identified proteins contain α/β serine hydrolase domains, and several are essential for parasite growth. One of the essential targets displays high homology to human monoacylglycerol lipase (MAGL) and is able to process lipid esters including a MAGL acylglyceride substrate. This Sal A target is inhibited by the anti-obesity drug Orlistat, which disrupts lipid metabolism and produces disorganized and stalled schizonts similar to Sal A. Resistance selections yielded parasites that showed only minor reductions in sensitivity and that acquired mutations in a protein linked to drug resistance in *Toxoplasma gondii*. This inability to evolve efficient resistance mechanisms combined with the non-essentiality of human homologs makes the serine hydrolases identified here promising antimalarial targets.

## INTRODUCTION

*Plasmodium falciparum* (*Pf*) is a protozoan parasite that causes the most severe form of human malaria. Malaria threatens 40% of the world’s population, resulting in an estimated 220 million cases and nearly 440,000 deaths annually (WHO, 2018). Due to a lack of effective vaccines against the parasite, malaria treatment and control efforts rely on the administration of antimalarial drugs (White et al., 2014). In recent years, artemisinin-based combination therapies (ACTs) have been the first-line antimalarial drugs and have contributed significantly to a reduction in the global malaria burden (Tu, 2011; White, 2008). However, emerging resistance to artemisinin and the partner drugs used in ACTs threatens to reverse the gains that have been made against this disease (Dondorp et al., 2017; van der Pluijm et al., 2019). Therefore, there is a continuing need to identify novel targets and pathways against which efficacious therapeutics can be developed, both to widen the scope of treatment and to overcome existing mechanisms of antimalarial drug resistance.

Natural products have evolved to encompass a broad spectrum of chemical and functional diversity and structural complexity, which enables them to target diverse biological macromolecules in a highly selective fashion. These features of natural products not only suggest that they may hold much promise for the development of therapeutic agents but also make them ideal starting points for the development of chemical probes (Harvey, 2000; Li and Vederas, 2009; Rodrigues et al., 2016; Thomford et al., 2018). Natural product-inspired chemical probes have been instrumental in identifying therapeutic targets and studying poorly understood biological processes and fundamental biology, yet the targets of most natural products remain unknown (Bottcher et al., 2010; Carlson, 2010; Kreuzer et al., 2015; Taunton et al., 1996). Salinipostin A (Sal A) is a natural product produced by the bacterium *Salinispora* sp. that was isolated from a marine sediment. Sal A displays potent activity against *Pf* parasites, with an EC_50_ of 50 nM and a selectivity index of >1,000 over a variety of mammalian cell lines (Schulze et al., 2015). In a previous effort to identify drug targets of Sal A, selection experiments were attempted but failed to generate resistant parasites. These results suggested that Sal A might target multiple proteins or essential pathways for parasite development, and that its potency is not easily compromised by single point mutations. The refractoriness of Sal A to parasite resistance mechanisms combined with its potent and selective activity, and lack of structural similarity to other antimalarials make Sal A an interesting chemical tool to identify druggable pathways in *Pf* parasites.

The α/β hydrolase superfamily is one of the largest groups of structurally related proteins that encompass a diverse range of catalytic activities. Members of this family are composed of largely parallel β-sheets flanked on their terminal ends by α-helices. The α/β hydrolase domain contains a conserved catalytic dyad or triad formed by a nucleophilic serine residue and an acid-base residue pair. The archetypal α/β hydrolases are esterases, which catalyze two-step nucleophilic substitution reactions to mediate hydrolysis of esters, but substrates for α/β hydrolases also include amides and thioesters (Carr and Ollis, 2009; Holmquist, 2000; Nardini and Dijkstra, 1999; Ollis et al., 1992). In recent years, α/β serine hydrolases have attracted considerable attention for their critical roles in metabolic processes in humans. These metabolic serine hydrolases have been implicated in neurotransmission, pain sensation, inflammation, cancer, and bacterial infection (Bachovchin and Cravatt, 2012; Long and Cravatt, 2011). The *Pf* genome encodes about 40 putative members of the serine hydrolase superfamily based on annotated sequence homologies (Aurrecoechea et al., 2009). However, most of these have not been functionally characterized. Natural product analogs of cyclipostins and cyclophostins, which contain the same reactive functional core electrophile as Sal A target human and mouse α/β serine hydrolases as well as α/β serine hydrolases involved in lipid metabolism in *Mycobacterium tuberculosis* (*M. tb.)* (Madani et al., 2019; Malla et al., 2011; Nguyen et al., 2017; Nguyen et al., 2018). These data suggest that α/β hydrolases are promising candidate targets for Sal A in *Pf*.

Activity-based protein profiling (ABPP) has emerged as a powerful and versatile chemical proteomic platform for characterizing the function of enzymes and for rapidly identifying novel targets (Chen et al., 2016; Cravatt et al., 2008; Spradlin et al., 2019; Wright and Sieber, 2016). Central to ABPP is the use of activity-based probes (ABPs), which covalently modify the active sites of enzymes. The bicyclic phosphotriester core in salinipostin A is hypothesized to form a covalent bond with target proteins upon nucleophilic attack by a catalytic serine or cysteine residue in the target. We therefore devised an analog of salinipostin A that contains an alkyne group at the terminus of its lipid tail, thereby enabling its use for affinity isolation of labeled target proteins. Herein, we describe the use of this probe to identify multiple essential α/β serine hydrolases including an essential parasite ortholog of human monoacylglycerol lipase (MAGL) that appears to be one of several targets. We also report that resistance appears to involve mutations in a separate determinant that in *Toxoplasma gondii* functions as a multidrug resistance mediator. Collectively, our data provide evidence that Sal A inhibits multiple essential serine hydrolases in *Pf*, adversely impacting lipid metabolism, and implicate a new set of metabolic targets for future antimalarial drug discovery and development efforts.

## RESULTS

### Salinipostin A targets human and parasite depalmitoylases

The Sal A molecule is made up of two parts, namely a reactive cyclic phosphate group that can act as an electrophile for serine and cysteine nucleophiles and a lipid tail that presumably mimics a native cellular lipid. This saturated 15 carbon lipid tail aligns with the 16 carbons in palmitic acid such that the reactive phosphorus is located directly at the site of the carbonyl used to form a thiol ester between palmitate and a cysteine on a palmitoylated protein (Couvertier et al., 2014) (Fig. 1A). We therefore reasoned that Sal A might act as a substrate mimetic inhibitor of the serine hydrolases that perform depalmitoylation reactions on proteins (Davda and Martin, 2014; Won et al., 2018). We conducted a competition experiment in which whole *Pf* parasites were incubated with Sal A prior to labeling with the broad-spectrum serine hydrolase activity-based probe fluorophosphonate-rhodamine (FP-Rho) (Simon and Cravatt, 2010). Several hydrolases were inhibited upon treatment with 1 μM Sal A (Fig. 1B). While palmitoylation is a ubiquitous and dynamic post-translational modification in *Pf* parasites (Corvi et al., 2012; Corvi and Turowski, 2019; Jones et al., 2012), there are no validated depalmitoylases in this parasite. Therefore, we tested Sal A in the related parasite pathogen, *Toxoplasma gondii*. To determine whether Sal A inhibits the *T. gondii* depalmitoylase PPT1 (TgPPT1), we treated wild-type (Δku80) and TgPPT1 knockout (ΔPPT1) parasites (Child et al., 2013) with Sal A and labeled parasite extracts with FP-Rho. These studies confirmed that Sal A was able to compete with FP-Rho for labeling of TgPPT1, as well as several other serine hydrolases in *T. gondii* (Fig. 1C). Competition studies in human cells (HEK293T) confirmed that Sal A inhibits the human depalmitoylases APT1 and APT2 in a dose-dependent manner, with increased potency against APT2 over APT1 (Fig. 1D). In addition, several other less abundantly labeled human hydrolases were also inhibited by Sal A pretreatment suggesting that Sal A likely has multiple serine hydrolase targets.

**Figure 1.**
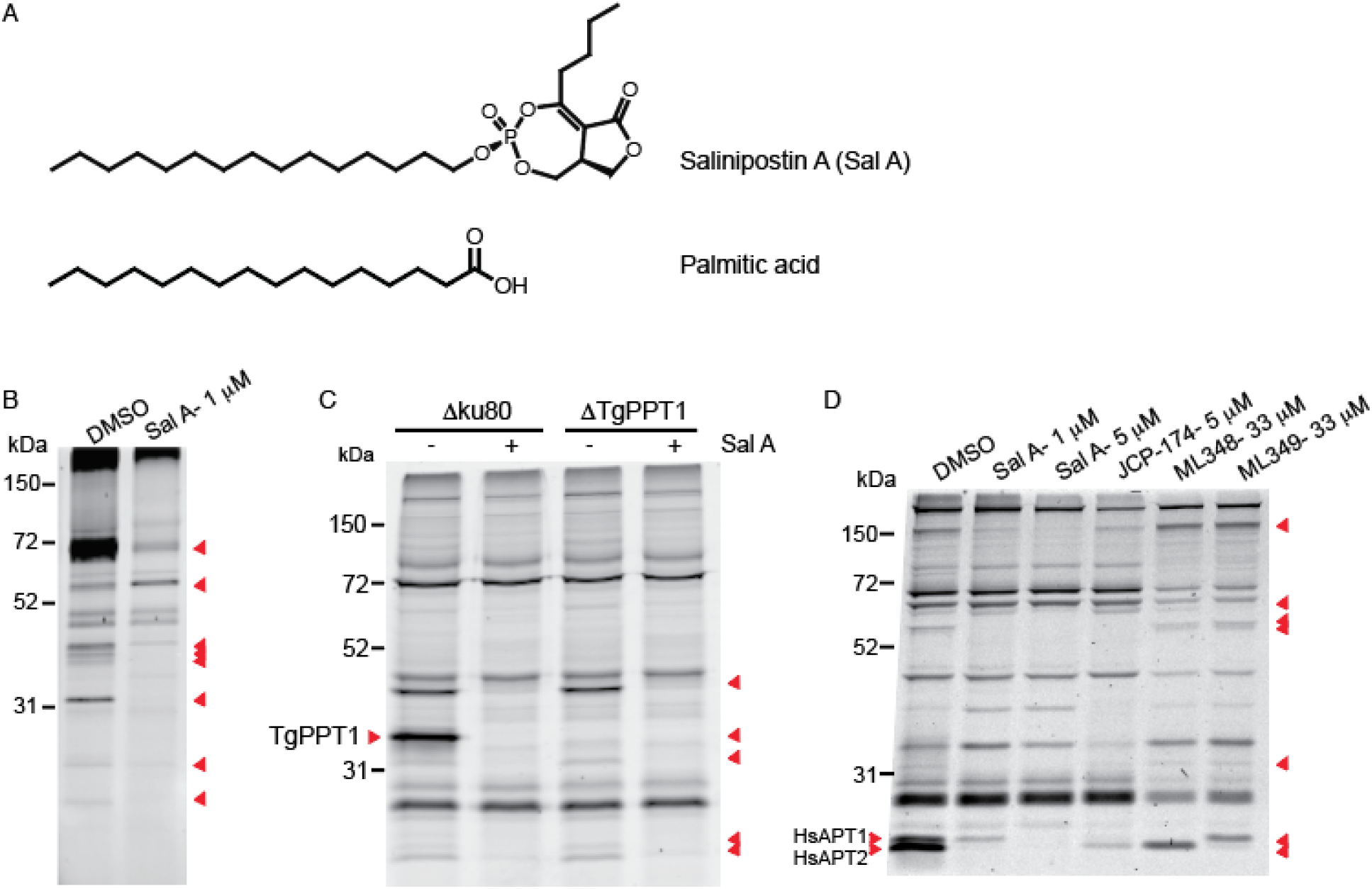
Effect of the natural product Salinipostin A on *Toxoplasma gondii* and human delpalmitoylases. (A) Chemical structure of salinipostin A (Sal A) and palmitic acid. (B-D) Competition assays with serine-hydrolase-reactive fluorophosphonate-rhodamine (FP-Rho) probe identifies multiple serine hydrolases as potential targets in (B) *P. falciparum* parasites, (C) *Toxoplasma gondii* Δku80 and ΔPPT1 parasites, (D) HEK293T cells with and without preincubation of Sal A.

### Synthesis of salinipostin A chemical probe

Since Sal A likely forms covalent bonds with its targets through reaction at the electrophile phosphorus (Fig. 2A), it was ideally suited to be converted into a probe that could be used to isolate and identify binding proteins in *Pf*. Sal A was originally isolated along with 10 analogs, which provided important structure-activity relationship (SAR) information for the design of the chemical probe. Analysis of these analogs revealed a significant drop in potency against parasites as the length of both the northern alkyl chain (C4>C3>C2) and the western alkyl chain (C15>C14>C13) decreased (Schulze et al., 2015). To minimize structural deviations from the natural product scaffold, we designed a chemical probe with a butyl northern alkyl chain and a terminal alkyne on the western alkyl chain (C16), thus introducing a single carbon and two degrees of unsaturation to the parent molecule. While total syntheses of similar natural products have been accomplished (Malla et al., 2011; Zhao et al., 2018), the bicyclic phosphotriester core of Sal A presents numerous synthetic challenges. We therefore opted for a semi-synthetic route in which the natural product Sal A was dealkylated at the western chain and the resulting phosphate diester was reacted with an electrophilic alkyne to form two diastereomers of the alkynylated natural product (Sal alk; Fig. 2B). Screening of these diastereomers against *Pf* parasites revealed that the isomer corresponding to the parent natural product retained full potency, whereas the other diastereomer showed a 10-fold drop in potency (Fig. S1). Additionally, we conducted 72 hour growth assays in the presence and absence of isopentenyl pyrophosphate (IPP), which is a product of apicoplast metabolism that is essential for growth during the asexual blood stage. An apicoplast-based killing mechanism can be reversed by supplementation of parasite culture medium with IPP (Yeh and DeRisi, 2011). The potency of both Sal A and Sal alk was unaffected by IPP, suggesting that these compounds do not target the apicoplast (Fig. S1).

**Figure 2.**
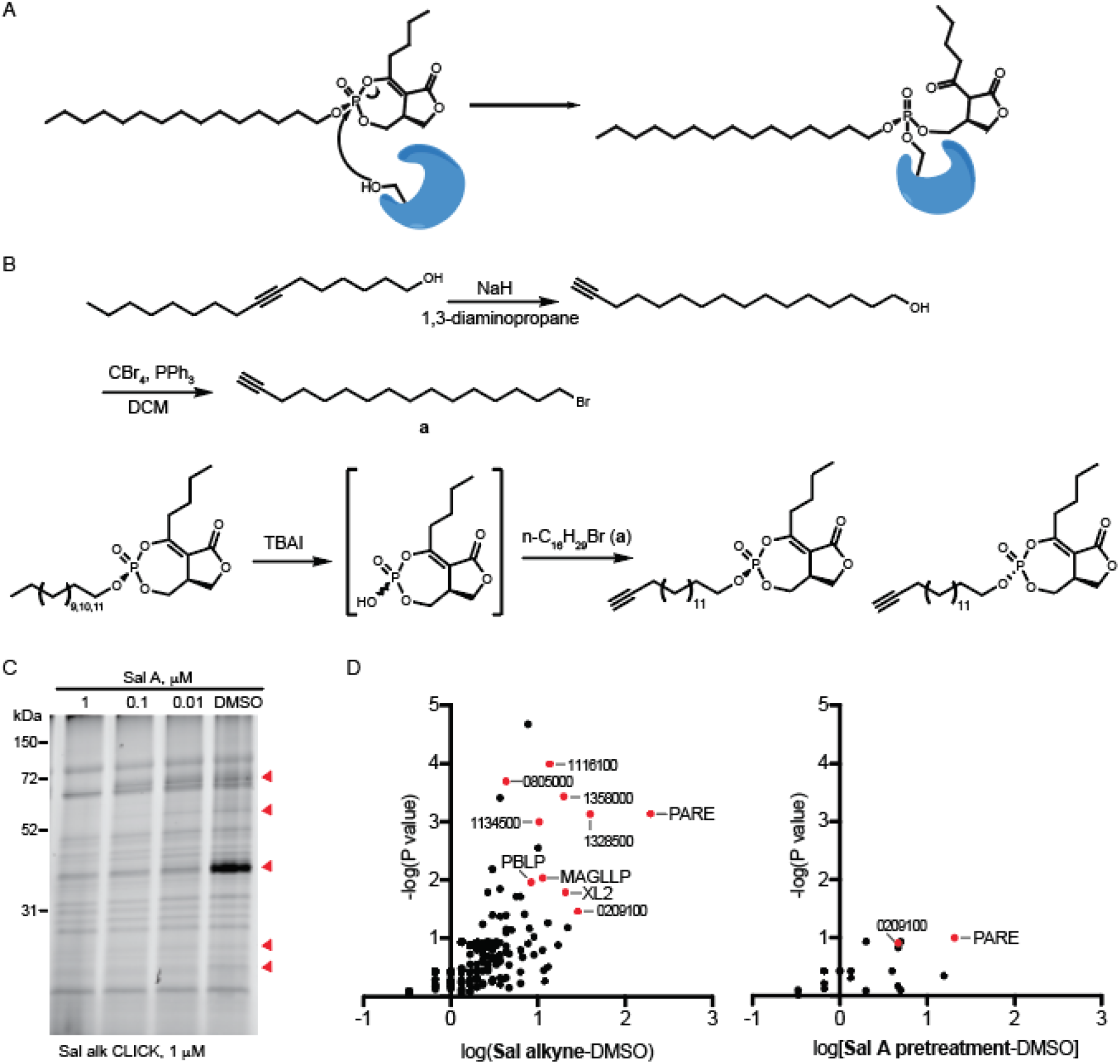
Semi-synthesis of an alkyne-labeled analog of Sal A and its application for chemical proteomics. (A) Proposed mode of action of Sal A. Sal A, which can form a covalent adduct with the nucleophilic serine catalytic residues at the active site of α/β hydrolase enzymes, rendering them inactive. (B) Semi-synthesis of Sal A alkyne. DCM, dichloromethane; TBAI, tetrabutylammonium iodide (see also Figure S1). (C) Sal alk labeling of proteins in *P. falciparum* with and without Sal A preincubation. After the Sal alk probe labeling, lysates of saponin-isolated parasites were prepared and labeled proteins were conjugated with TAMRA-azide using CLICK reaction (see also Figure S2). (D) Volcano plots of Sal A alkyne targets in *P. falciparum*. X axis shows the logarithm values of difference in spectral counts of protein between Sal alkyne-treated and DMSO vehicle-treated parasites (left) and Sal A-pretreated (competition) and DMSO vehicle-treated parasites (right). The statistical significance was determined without correcting for multiple comparisons in triplicates and the Y value in the volcano plot is minus one times the logarithm of the P value. The proteins with high abundance and competition by the parent compound Sal A are highlighted in red (see also Table 1 and Table S1 for identified proteins in red).

**Table 1.** Sal A target proteins identified in *P. falciparum*

| Protein ID | Description | MW (kDa) | Essential in *P. falciparum* piggyBac screen | Spectral counts |
| --- | --- | --- | --- | --- |
| PF3D7_0709700 | lysophospholipase (PfPARE) | 42.4 | No | 195.0 |
| PF3D7_1328500 | $\alpha/\beta$ hydrolase | 115.9 | No | 39.7 |
| PF3D7_0209100 | phospholipase A2 | 78.3 | No | 28.7 |
| PF3D7_1001600 | exported lipase 2 (PfXL2) | 88.6 | Yes | 20.7 |
| PF3D7_1358000 | patatin-like phospholipase | 238.2 | No | 19.7 |
| PF3D7_1116100 | serine esterase | 217.1 | No | 13.7 |
| PF3D7_1038900 | esterase, lysophospholipase | 41.8 | Yes | 11.3 |
| PF3D7_1134500 | $\alpha/\beta$ hydrolase | 210.5 | Yes | 10.3 |
| PF3D7_0818600 | BEM46-like protein (PBLP) | 34.9 | Yes | 8.3 |
| PF3D7_0805000 | $\alpha/\beta$ hydrolase | 28.5 | No | 4.3 |

### Salinipostin A targets multiple serine hydrolases in *P. falciparum* parasites

Having confirmed that Sal alk retained activity equivalent to the parent natural product, we conducted a competition experiments in which parasites were pre-incubated with Sal A for 2 hours prior to Sal alk labeling. CLICK reaction with TAMRA-azide confirmed that Sal alk labeled multiple proteins in *Pf* and that Sal A competed for Sal alk labeling of several of those proteins in a dose-dependent manner (Fig. 2C). Therefore, we next used Sal alk as a probe to affinity purify targets of the natural product using a standard ABPP method. For these studies, we pretreated *Pf* parasite cultures with either Sal A or a vehicle control and subsequently labeled cultures with Sal alk. Samples were lysed and labeled proteins were conjugated to biotin-azide using CLICK chemistry. Avidin affinity chromatography was performed to enrich for biotin-azide labeled proteins. The samples were then subjected to tryptic digestion and tandem mass spectrometry analysis (Table S1). Analysis of these data identified 10 putative Sal A targets that were enriched in Sal alk-treated samples and that were correspondingly reduced in Sal A-pretreated samples (Fig. 2D, Table 1, and Table S1). All ten putative target proteins have α/β serine hydrolase domains with either a conserved Ser-His-Asp catalytic triad or a Ser-Asp dyad. Of these, five are annotated as lipases with the characteristic GXSXG motif (http://plasmodb.org/plasmo) (Aurrecoechea et al., 2009). Based on a recent piggyBac transposon saturation mutagenesis study (Balu et al., 2009; Balu et al., 2005; Bushell et al., 2017; Zhang et al., 2018), four of the identified targets are likely to be essential in *Pf*.

One of the proteins identified as a putative Sal A target, PF3D7_1001600 was recently annotated as exported lipase 2 (PfXL2), one of the *Plasmodium* α/β hydrolases containing the export element (PEXEL) motif that is essential for delivery of most exported proteins (Hiller et al., 2004; Marti et al., 2004). Studies with trophozoite/early schizont-stage parasites expressing a C-terminal GFP fusion of PfXL2 revealed that the protein is exported into the erythrocyte cytosol of the infected RBCs (Spillman et al., 2016). Nonetheless, the physiological function of PfXL2 remains unknown, although it is considered to be essential for parasite growth (Bushell et al., 2017; Zhang et al., 2018).

We also identified a *Plasmodium* BEM46-like protein (PBLP, PF3D7_0818600) as another potential target of Sal A. A recent study of PBLP in the rodent malaria parasite *Plasmodium yoelii* showed that it has an important role in invasion, with deletion of PBLP affecting merozoite formation, sporozoite maturation, and infectivity (Groat-Carmona et al., 2015). The bud emergence (BEM) 46 proteins are an evolutionarily conserved class of α/β hydrolases that carry N-terminal signal peptides that can act as transmembrane domains. BEM46 homologues have been implicated in signal transduction and cell polarity (Kumar et al., 2013; Mercker et al., 2009). All *Plasmodium* species appear to express a PBLP orthologue, which might suggest a conserved function. However, the exact function of PBLP is still unclear.

Based on total spectral counts, the most abundantly labeled potential Sal A target was a putative lysophospholipase (PF3D7_0709700). This protein was recently annotated as PfPARE (*P. falciparum* Prodrug Activation and Resistance Esterase), an esterase that is responsible for activating some antimalarial prodrug compounds (Istvan et al., 2017). To test whether PfPARE could be a Sal A target, we obtained *Pf* lines expressing both wild-type (WT-GFP) and active site mutant (S179T-GFP) PfPARE-GFP fusions replacing the native gene locus. Treatment of these parasites with Sal alk revealed that labeling of PfPARE that was dependent on the catalytic serine (Fig. S2A). However, parasites expressing the WT or catalytic dead PfPARE were equally sensitive to Sal A treatment (Fig. S2B). These results, combined with the predicted lack of essentiality of PfPARE, suggest that it most likely binds Sal A because of its general drug binding properties and that it may not be a relevant target for the compound’s parasiticidal action.

### Salinipostin A inhibits a monoacylglycerol lipase (MAGL)-like protein in *P. falciparum* (PfMAGLLP)

To gain further insight into potential functions of the serine hydrolases that we identified as putative Sal A targets, we performed a sequence homology search and found that three of the ten proteins (PfPARE, PfXL2, and PF3D7_1038900) share sequence similarity with human monoacylglycerol lipase (MAGL; Fig. 3A and Fig. S3). It is possible that these proteins all have MAGL-like properties. Unfortunately, we were unable to express and purify PfXL2. A recent study on PfPARE confirmed its esterase activity and broad substrate specificity. However, the catalytic activity of this enzyme is dispensable for asexual blood stage growth, as active site mutant parasites (S179T) displayed no growth defect (Istvan et al., 2017). The third protein with sequence similarity to human MAGL, PF3D7_1038900, is predicted to encode a hydrolase closely related to PfPARE, with 53% sequence identity over 338 residues. This previously uncharacterized enzyme, referred to henceforth as PfMAGLLP is predicted to have a common structural core α/β hydrolase fold composed of largely parallel β-sheets flanked by α-helices, and it contains the conserved catalytic Ser-His-Asp triad of serine hydrolases (Fig. 3B and Fig. S4A). Furthermore, the cysteine residue adjacent to the active site that is important for ligand recognition and enzyme activity of human MAGL is conserved in the *Plasmodium* enzyme (Karlsson et al., 1997; Saario et al., 2005).

**Figure 3.**
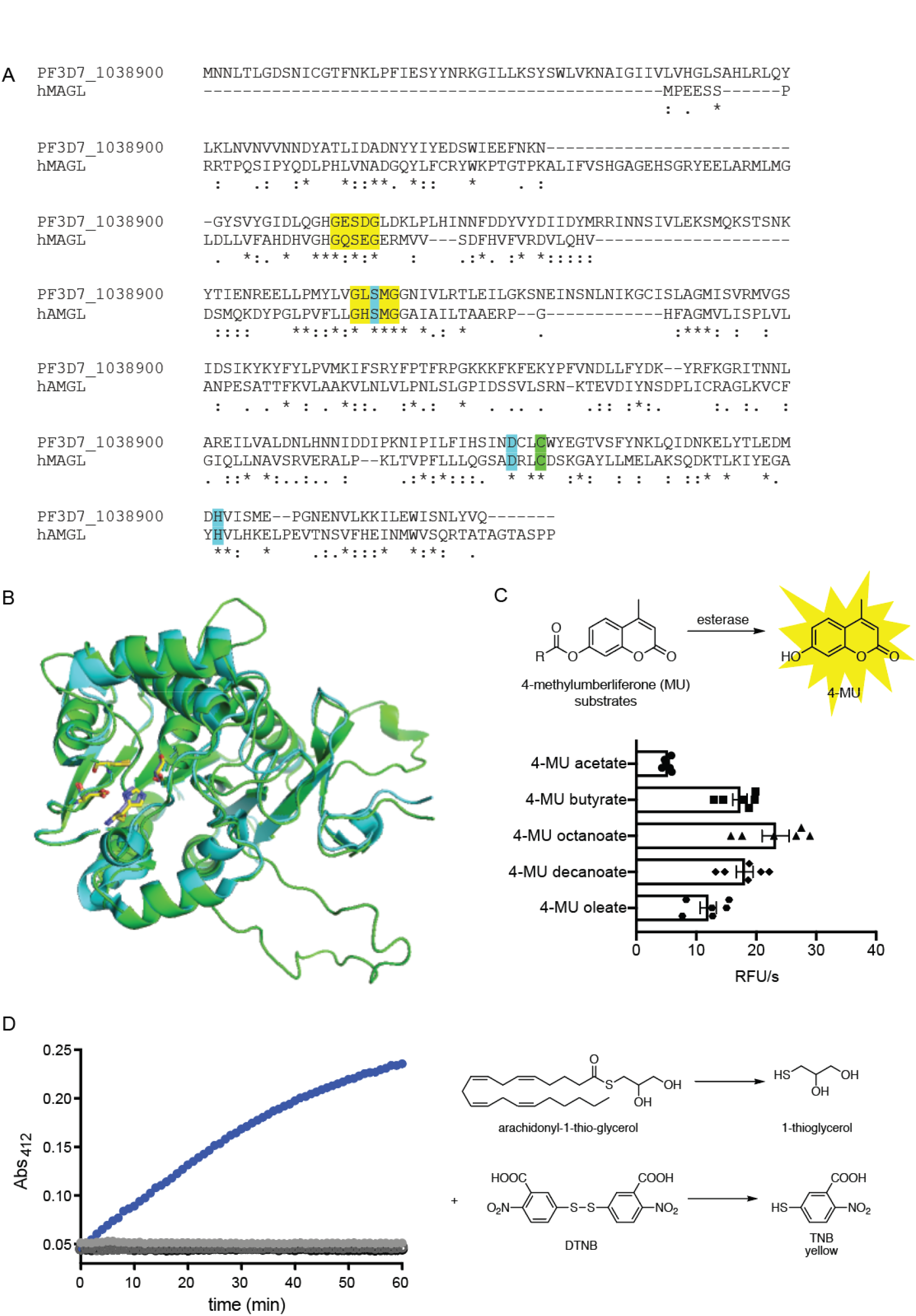
Sequence, structural alignment and activity of the serine hydrolase target of Sal A, PF3D7_1038900 (PfMAGLLP). (A) Sequence alignment of PfMAGLLP and hMAGL using Clustal W2. Identical and highly similar residues are indicated with (*) or (:), respectively. Sequence identity is 17% and FFAS (fold and function assignment system) score is −74.8. The conserved G-X-S-X-G motif and residues of the catalytic triad are highlighted in yellow and cyan, respectively. The cysteine adjacent to the active site is highlighted in green (see also Figure S3). (B) Predicted structure of PfMAGLLP (green) superimposed with human MAGL structure (PDB: 3jw8, cyan). Catalytic triads are shown in yellow for PfMAGLLP and purple for hMAGL (see also Figure S4A). (C) Processing of 4-methylumbelliferyl (4-MU)-based fluorogenic substrates by recombinant PfMAGLLP. 10 μM of substrates were incubated with 5 nM of rPfMAGLLP and initial cleavage rates (relative fluorescence units/sec) were measured. Data were derived from three independent experiments performed in duplicate and were calculated as non-linear regressions using GraphPad Prism with means ± s.d (see also Figure S4B). (D) Hydrolysis of arachidonyl-1-thio-glycerol by rPfMAGLLP. Hydrolysis of the thioester bond by the enzyme generates a free thiol that reacts with 5,5’-dithiobis-(2-nitrobenzoic acid) (DTNB) resulting in a yellow product thionitrobenzoic acid (TNB) with an absorbance of 412 nm (blue). In the absence of either enzyme or substrate, there was no absorbance of TNB (Baragana et al.).

*In vitro*, human MAGL is capable of hydrolyzing a variety of monoglycerides into free fatty acids and glycerol. This enzyme is responsible for the hydrolytic deactivation of the endocannabinoid 2-arachidonoyl-sn-glycerol (2-AG) and has been implicated in diverse pathophysiological processes, including pain, inflammation, and neuroprotection (Labar et al., 2010; Long et al., 2009a; Long et al., 2009b). To determine whether the PfMAGLLP is structurally and functionally related to human MAGL, we recombinantly expressed and purified the protein and evaluated its ability to process lipid substrates. PfMAGLLP was able to process various fluorogenic lipid ester substrates bearing different chain lengths including butyrate, octanoate, and monosaturated oleate (Fig. 3C). Among the substrates tested, 4-methylumbelliferyl (4-MU) octanoate proved to be an optimal substrate with the highest k_cat_/K_m_ value (Fig. S4B). PfMAGLLP also efficiently hydrolyzed arachidonoyl-1-thio-glycerol, a close analog of the verified MAGL substrate 2-AG. This substrate produces 1-thioglycerol, which can be subsequently detected by the release of a yellow thiolate ion (TNB) upon reaction with 5,5′-dithiobis(2-dinitrobenzoic acid) (DTNB, Ellman’s reagent(Ellman, 1959); Fig. 3D).

With an active recombinant enzyme and corresponding fluorogenic substrate in hand, we were able to initiate screens for inhibitors of PfMAGLLP. We first tested reported human MAGL inhibitors containing the carbamate electrophile, including KML29, MJN110, JW651, and JZL 184 (Chang et al., 2013; Chang et al., 2012; Niphakis et al., 2013). At 1 μM concentrations, MLN110, JW651, and KML29 inhibited the esterase activity of PfMAGLLP by more than 90% while JZL 184 inhibited activity by 75% (Fig. S5A). Dose-response assays showed a broad spectrum of potencies for the four MAGL inhibitors, with IC_50_s ranging from 2.6 nM for JW651 to 163 nM for KML29 (Fig. 4A). JW651 had similar potency against recombinant PfMAGLLP as did Sal A, thereby making the compound a useful tool to further explore the function of this enzyme in malaria parasites.

**Figure 4.**
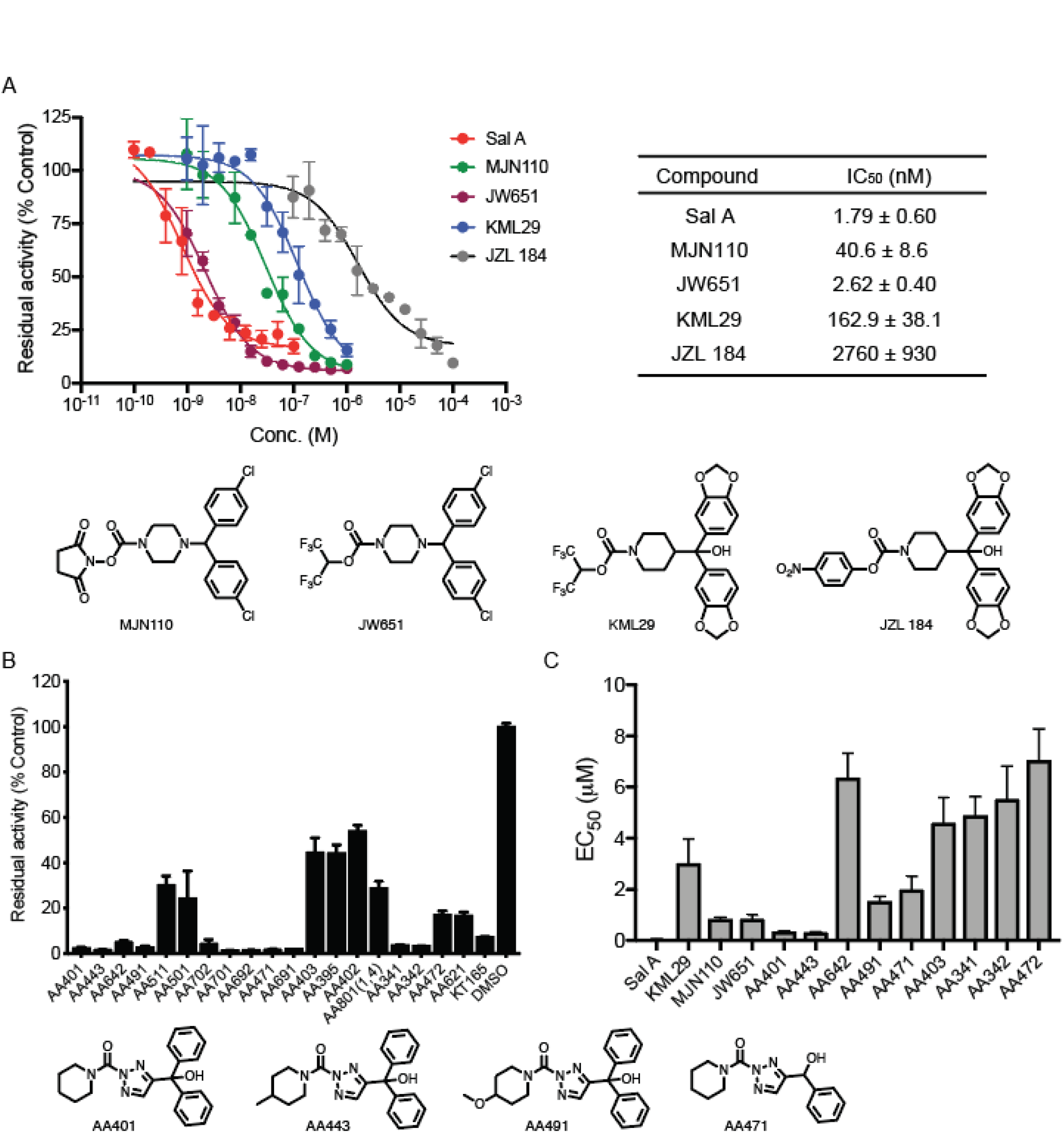
Effects of Human MAGL inhibitors and triazole ureas identified from screening on recombinant PfMAGLLP and parasite growth in culture. (A) Inhibition of PfMAGLLP as determined by initial cleavage rates (relative fluorescence units/sec) by known human MAGL inhibitors (structures shown below, see also Figure S5A). Reactions contained 5 nM rPfMAGLLP and 10 μM substrate 4-methylumbelliferyl octanoate substrate. Data were derived from three independent experiments performed in duplicate and were calculated as non-linear regressions using GraphPad Prism. IC_50_ values are listed (Mean ± s.d.). (B) Inhibition of rPfMAGLLP with a small library of triazole urea containing compounds as determined by normalized initial cleavage rates to DMSO control. The top 20 selected compounds were screened at 100 nM concentration (see also Figures S5B-D). (C) EC_50_ values of PfMAGLLP inhibitors in 72 h treatment of *P. falciparum* W2 parasites, beginning as synchronized ring stages with error bars, s.d. (n = 6 parasite cultures from two independent experiments with triplicates). Structures of representative compounds are shown.

We next performed the screening of a small library of 154 serine reactive triazole-urea containing compounds and identified 31 hits with greater than 90% inhibition of recombinant PfMAGLLP at 1 uM concentrations (Fig. S5B). Further screening of the top 20 most potent hits at lower concentrations identified multiple compounds with strong inhibitory activity in the nM range (Fig. 4B). These compounds were also screened for their ability to compete for active site labeling of PfMAGLLP and human MAGL with the general serine hydrolase probe FP-Rho in order to identify compounds with the highest potency as well as the greatest selectivity for the parasite enzyme (Fig. S5C).

To determine whether the potency of inhibition of PfMAGLLP for a given compound correlated with its *in vitro* potency against parasites, we performed dose-response parasite growth inhibition assays with the human MAGL inhibitors and the top triazole urea hits (Fig. 4C). Assays with W2 parasites revealed that the anti-parasitic potencies for this set of compounds largely correlated with the potencies against the recombinant PfMAGLLP enzyme. The most pronounced exception was the compound JW651, which had virtually identical potency as Sal A against recombinant PfMAGLLP, but showed a 16-fold decrease potency as compared with Sal A against parasites (EC_50_ value of 839 nM vs 50 nM for Sal A). This discrepancy could be due to reduced cell permeability of JW651 or could be the result of Sal A targeting multiple serine hydrolases in the parasite. In further support of the hypothesis that Sal A functions by targeting multiple serine hydrolases, we performed dose-response assays with parasites overexpressing PfMAGLLP from an exogenously transfected plasmid. These parasites showed no apparent shift in Sal A EC_50_ values as compared with wild-type parasites (Fig. S6). Similar results were obtained for two of the other putative Sal A targets (PF3D7_0808500 and PF3D7_0818600) in that no shift in EC_50_ was observed in parasites exogenously overexpressing these genes. These results further support the hypothesis that Sal A induces parasite killing by targeting multiple serine hydrolases.

### Orlistat is a potent inhibitor of PfMAGLLP

Orlistat (Fig. 5A), the anti-obesity drug that irreversibly inhibits gastric and pancreatic lipases that hydrolyze triacylglycerols (TAG) to fatty acids, was also recently found to inhibit the thioesterase domain of fatty acid synthase (FAS) (Pemble et al., 2007; Weibel et al., 1987). Additionally, recent studies of Orlistat in *Pf* suggest that the compound functions by disrupting parasite neutral lipid metabolism pathways (Gulati et al., 2015). However, the specific targets of Orlistat in *Pf* have not been determined.

**Figure 5.**
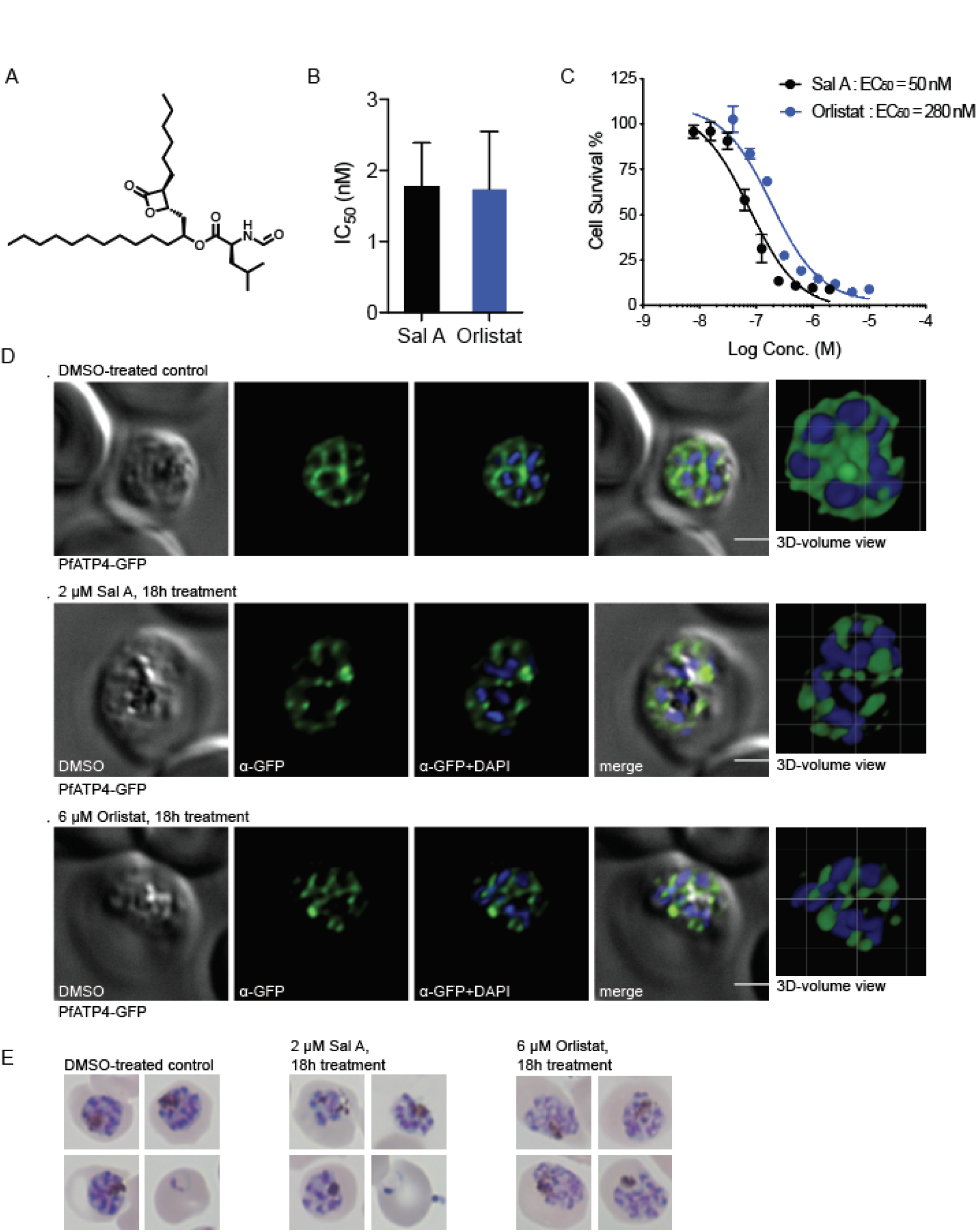
Effects of the anti-obesity drug Orlistat on parasite growth compared to Sal A.

(A) Chemical structure of Orlistat. (B) IC_50_ values of Sal A and Orlistat for inhibition of 4-methylumbelliferyl octanoate processing by rPfMAGLLP. (C) EC_50_ values of Sal A and Orlistat in 72 h treatment of *P. falciparum* W2 parasites, beginning as synchronized ring stages with error bars, s.d. (D) Plasma membrane ingression around developing daughter cells visualized using the plasma membrane marker PfATP4-GFP (McNamara et al., 2013). Synchronized trophozoite-stage parasites were treated at 24-30 hpi with DMSO (vehicle control), 2 μM Sal A or 6 μM Orlistat and imaged 18 h later. Control parasites had organized and fully enclosed developing merozoites, whereas plasma membrane ingression was perturbed by Sal A or Orlistat treatment. Nuclei were stained with Hoechst 33342. (E) Giemsa-stained parasites exposed to the same treatments as in (D).

We tested Orlistat for its ability to inhibit recombinant PfMAGLLP and found that it was equipotent to Sal A, suggesting that the enzyme is a likely target (Fig. 5B). We next performed dose-response assays with Sal A and Orlistat and found that Orlistat is less potent at killing parasites than Sal A (EC_50_ value of 280 nM vs 50 nM for Sal A; Fig. 5C). To further explore the cellular effect of Sal A on asexual blood stage parasites, we took advantage of a *Pf* reporter line that expressed the plasma membrane protein PfATP4 fused to GFP. Fluoresence-microscopy imaging of mature schizonts expressing PfATP-GFP reporter and treated for 18h with the DMSO vehicle control reveals invagination of the plasma membrane around the individual merozoites during the final stages of cytokinesis and preparation for parasite egress from the infected RBC (Fig. 5D), as previously reported (Rottmann et al., 2010). In contrast, parasites treated for 18 h with 2 μM Sal A or 6 μM Orlistat showed disorganized plasma membranes that failed to properly surround the merozoites, indicating that both compounds prevented parasites from completing cytokinesis and egressing from the infected RBCs (Fig. 5D). These data recall a similar cellular block observed with the KAE609 inhibitor that targets the *Pf* phospholipid 4-kinase PfPI4K (McNamara et al., 2013).

Based on our recent results with Orlistat, which showed that the drug functions by blocking key lipid metabolism pathways resulting in accumulation of triacylglycerides (TAGs) (Gulati et al., 2015), we wanted to determine whether similar accumulations of lipid species could be detected in parasites treated with other putative PfMAGLLP inhibitors. We thus performed metabolomic studies on JW651-treated parasites to measure changes in key lipid metabolites. JW651 treatment resulted in an accumulation of monoglycerides, including palmitoyl and oleoyl glycerols, with a greater than two-fold increase in these products compared to DMSO-treated parasites (Fig. S7). Together, these results suggest that PfMAGLLP has a role in processing these lipid esters, and that Orlistat and JW651 block lipid processing, resulting in schizont arrest and parasite death. In further support of the essential role of PfMAGLLP, attempts to generate a conditional knockout of PfMAGLLP have not been successful.

### *In vitro* evolution of resistance to Sal A

Prior attempts at selecting for resistance to Sal A *in vitro* were unsuccessful at generating Sal A-resistant parasites (Schulze et al., 2015). These studies, which subjected 10^10^ parasites to 3 to 7-fold IC_50_ concentrations, in three separate experiments, yielded no resistant parasites. These selections were performed using W2 wild-type parasites, and the exposure to low, sublethal concentrations of Sal A (ranging from 160 to 360 nM) allowed a small subset of the parasite population to recrudesce without any demonstrable decrease in sensitivity. To improve our odds of selecting for resistance to Sal A, we made use of a parasite line that has a mutation in DNA polymerase delta (referred to herein as Dd2_Polδ). This polymorphism ablates the polymerase’s proofreading (exonuclease) activity, thereby increasing the overall DNA mutation rate (Honma et al., 2016). We performed *in vitro* resistance selections with Dd2_Polδ and Sal alk in duplicate flasks, each containing 1×10^9^ parasites. Parasites were exposed to 2.5 μM Sal alk (~2×the IC_90_ concentration of Sal alk against the Dd2_Polδ line) over the course of ten days, after which drug pressure was removed and the cultures were continued in regular media. Recrudescence was observed microscopically in both flasks at day 17 of the selection. Selected cultures displayed a small but significant shift in IC_50_ (50% growth inhibitory concentration) when assayed in bulk. Clones from both flasks were obtained by limiting dilution, and six (three from each flask) were selected for whole-genome sequencing (WGS) analysis. Across the six Sal A-selected clones sequenced, single-nucleotide polymorphisms (SNPs) were identified in 44 genes.

Among these genes, only two showed mutations in clones from both flasks. None of the mutations were in genes coding for α/β hydrolases, rather, the sole gene that was mutated in all six clones was PF3D7_1324400, which is annotated as a putative PRELI-domain containing protein of unknown function. Five of the six clones harbored a single point mutation in the coding sequence (A20V in clones Fl2D7 and Fl2G3 from flask 2, or P102L in Fl2A2, and T145I from the identical clones Fl1C12 and Fl1E12 from flask 1; Table S2). The sixth clone, Fl1A9, had a point mutation in intron 1 that has an unknown impact on expression levels. The second frequently mutated gene was PF3D7_1235200, encoding a V-type H^+^-translocating pyrophosphatase, which showed a S725Y mutation in the identical clones Fl1C12 and Fl1E12 and an E507K mutation in the clone Fl2D7. The absence of mutations in this gene in the other Sal A-resistant clones leads us to suspect that this gene is at best only an occasional contributor to the resistance phenotype and is not the primary determinant. Other mutations arising infrequently are thought to be stochastic events related to the hyper-mutability observed in this DNA_Polδ mutant line.

Drug susceptibility assays with these clones and the parental Dd2_Polδ line showed significant 2 to 9-fold increases in IC_50_ values for the resistant clones (Table S2), with dose-response data showing decreased susceptibility across multiple concentrations (Fig. 6A). No significant cross resistance was observed to Orlistat, known to inhibit the hydrolysis of triacylglycerols in *P. falciparum* (Gulati et al., 2015) (Fig. 6B). We also found no cross resistance to JW651 (Table S2), suggesting different specificities in the modes of action of Sal A and these two other lipid-targeting antimalarials.

**Figure 6.**
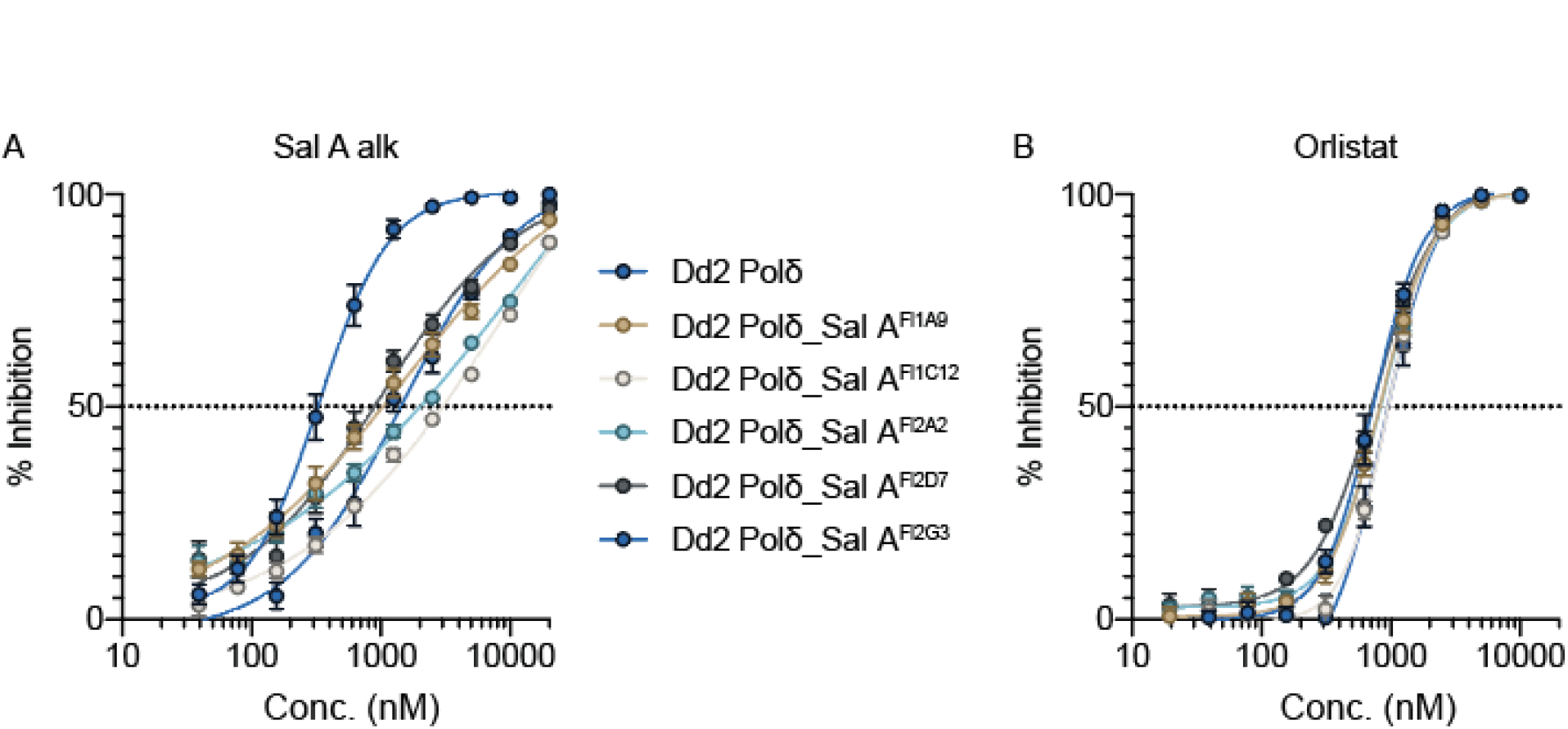
Parasites selected for Sal A resistance have a ten-fold reduction in sensitivity to Sal A but not to Orlistat. (A, B) Dose-response curves for Sal A-selected parasite clones and the corresponding parental line (Dd2 Polδ) tested against (A) Sal alk or (B) the lipid metabolism inhibitor Orlistat.

## DISCUSSION

Lipid-dependent processes, including the formation of a vacuolar system and the development of daughter cells, play crucial roles in the *Plasmodium* life cycle, and are fueled both by *de novo* synthesis and by acquisition of lipids from the host cell (Ben Mamoun et al., 2010; Vial et al., 2003). Yet the specific enzyme(s) responsible for lipid turnover during the replication and invasion stages of malaria parasites remain poorly understood. Here we show that a natural product, Sal A, results in stage-specific inhibition of parasite development by targeting multiple α/β serine hydrolases. This stage-specificity profile is similar to that observed for the anti-obesity drug Orlistat, previously shown to block lipid metabolism pathways in *Plasmodium* parasites (Gulati et al., 2015). One of the primary Sal A targets identified here, PfMAGLLP, demonstrated esterase activity for lipid substrates including monoacylglycerols and was also potently inhibited by Orlistat. Biochemical characterization and inhibitor screening of this hydrolase led to the identification of small molecule inhibitors that similarly blocked parasite development and led to accumulation of acylglyceride metabolites. Understanding the roles of *Plasmodium* α/β serine hydrolases in host and parasite lipid metabolic networks will likely open up new avenues to discover and develop antimalarials with novel mechanisms of action.

The original studies that identified Sal A as a potent natural product antimalarial postulated that it functioned by disrupting key parasite signaling pathways, and efforts were subsequently made to identify direct targets of the molecule using drug resistance screening (Schulze et al., 2015). These efforts failed to produce resistant mutants and the authors suggested that the compound might kill parasites through pleotropic effects on multiple targets, making it unlikely that resistant mutants could evolve through single nucleotide polymorphisms. In support of this multi-target hypothesis, studies of the related natural products, cyclipostins and cyclophostins, which contain the same type of phosphate electrophile found in salinipostins, have revealed potent inhibitory activity against multiple classes of serine lipases and esterases. These compounds are also potent against *Mycobacterium tuberculosis* (*M. tb.*), and activity-based protein profiling (ABPP) efforts have identified a number of serine hydrolase targets in that species (Madani et al., 2019; Malla et al., 2011; Nguyen et al., 2017; Nguyen et al., 2018). A recent review of the cyclipostins and related compounds suggested that, given the broad-spectrum but class-specific reactivity of the cyclipostins, Sal A might function through inhibition of multiple serine hydrolase targets in *Plasmodium* (Spilling, 2019). We initially hypothesized that Sal A might target serine hydrolases responsible for removing palmitate from S-palmitoylated proteins, given its structural similarity to palmitic acid. Indeed, we found that Sal A was able to target *Toxoplasma* depalmitolyase (TgPPT1) and induce a parasite growth and invasion arrest. However, several other serine hydrolases were also effectively targeted by the compound, supporting a generalized mode of action that involves inhibition of multiple enzymes with likely similar catalytic mechanisms and substrates.

The field of chemical biology has evolved over the past few decades, but a common theme has been the use of small molecules to dissect biological pathways. While there are examples of small molecules that can be used to perturb specific biological pathways by binding to a single protein target, in many cases, small molecules have effects that are mediated by their ability to interact with multiple protein targets. On the one hand, these types of pleotropic effects make mechanistic studies of small molecules much more challenging, on the other, they make many small molecules useful tools for identifying related proteins that function together in important biological pathways. For Sal A, the results described herein point to a mechanism of action involving disruption of multiple essential serine hydrolases involved in lipid metabolism. While we were unable to identify which of these hydrolase targets is most integral to the compounds antimalarial activity, likely because parasite killing results from the combined inhibition of several targets, our data identify a number of previously poorly characterized enzymes that, collectively, appear to be essential for parasite survival.

We focused our attention on the three serine hydrolases identified as putative targets of Sal A that have been considered essential in *Pf* based on a recent whole-genome saturation mutagenesis study. Homology searches and predicted structural analyses of these enzymes identified one, PF3D7_1038900, as a likely MAGL protein, due to its sequence homology to the human MAGL involved in processing of acylglycerides and other lipid metabolites. Recombinant expression of this protein, PfMAGLLP, resulted in an active enzyme that was able to process both simple lipid esters as well as a close homolog of the native human MAGL substrate 2-arachidonoyl-glycerol (2-AG). Both Sal A and the anti-obesity drug Orlistat were potent inhibitors of this enzyme. We were also able to identify several novel inhibitors of PfMAGLLP and show that human MAGL inhibitors also effectively block its activity. While we cannot definitively confirm that inhibition of this target by Sal A is the primary cause of parasite killing, we found that the ability of similar compounds to inhibit PfMAGLLP generally correlated with their ability to kill parasites. Furthermore, we observed that a human MAGL inhibitor that is also a potent inhibitor of PfMAGLLP was able to induce accumulation of monoacylglycerides in treated parasites, again linking this enzyme to lipid metabolism in the parasite.

Our efforts to confirm specific serine hydrolases as phenotypically relevant targets of Sal A through drug selection studies resulted in the isolation of parasite clones with moderately diminished sensitivity to the drug. Specifically, we were able to isolate six clones from a hypermutating Dd2 clone that showed reproducible reduction in EC_50_ values of approximately 2 to 9-fold as compared with the parental line, and performed WGS on those clones. These data identified 44 genes with point mutations that resulted in either truncation or mutation of predicted protein products. While this number of mutations is high, the vast majority were found in only one or two of the sequenced clones. Only two genes showed multiple mutations in three or more of the clones and both might thus contribute to general modes of drug resistance. The most likely dominant mediator of the observed reduced sensitivity effect was a gene coding for a PRELI-domain containing protein of unknown function, for which mutations were found in all six clones obtained from both independent flasks. While the function of this protein in *Plasmodium* is unclear, a homolog from the related parasite pathogen *Toxoplasma gondii*, TgPREILD, was recently found to be a mitochondrial protein associated with multidrug resistance (Jeffers et al., 2017). Furthermore, PRELI domain-containing proteins in humans have been implicated in transfer of phospholipids to the inner mitochondrial membrane (Potting et al., 2013). Together these results suggest that parasite killing effects of Sal A are pleiotropic, and that point mutations in individual targets are not sufficient to generate resistance. Rather, resistance appears to occur via activation of a general drug-resistance pathway in the parasite. These results, combined with the fact that resistance to Sal A is mild and was difficult to generate, suggest that compounds with broad-spectrum activity against α/β hydrolases constitute exciting leads to develop novel antimalarials that can treat existing multidrug-resistant malarial infections.

## SIGNIFICANCE

Natural products can be powerful tools to identify novel biological pathways and targets in diverse organisms. Here we describe functional studies of the marine natural product Salinipostin A, a compound that was discovered as a highly potent and selective anti-malarial agent with a poorly defined mechanism of action. Using a semi-synthesis method, we generated a suitably tagged analog of Sal A for target identification studies. Using this probe, we identified a suite of largely uncharacterized serine hydrolases in *P. falciparum* parasites. Several of these enzymes are essential to parasite growth and are likely to be involved in diverse aspects of lipid metabolism and signaling. Our results suggest that the parasite killing effects of Sal A are likely the result of inhibition of multiple serine hydrolases. This mode of action makes generation of resistance through mutations in direct targets difficult. In fact, we find weak resistance to Sal A was mediated by mutations in a protein linked to drug resistance in a related parasite rather than by mutations in target genes. In addition, we found that the anti-obesity drug Orlistat induces a phenotypic block in parasite growth that is highly similar to that of Sal A. This drug, which has already been shown to block processing of host derived lipids, previously had no confirmed targets. At least one of the targets of Sal A is a likely homolog of human monoacylglycerol lipase (MAGL) and is also potently inhibited by Orlistat, thus liking it to essential lipid metabolism pathways in the parasite. Given the lack of essentiality of many of the human homologs of the identified serine hydrolases coupled with their critical function in parasites, our studies with Sal A have identified a number of poorly characterized enzymes that are attractive targets for new generation anti-malarial agents.

## METHODS

### *P. falciparum* replication assays

*P. falciparum* W2 cultures were maintained, synchronized, and lysed as previously described (Straimer et al., 2017). Parasite cultures were grown in human erythrocytes (no blood type discrimination) purchased from the Stanford Blood Center (approved by Stanford University). Ring stage *P. falciparum* culture at 1% parasitemia and 0.5% hematocrit was added to 96-well plates spotted with compounds. The *P. falciparum* culture was incubated with each compound for 72 h. After incubation, the culture was fixed in a final concentration of 1% paraformaldehyde (in PBS) for 30 min at room temperature. The nuclei stain YOYO-1 was then added to a final concentration of 50 nM and incubated overnight at 4°C. Parasite replication was monitored by observation of a YOYO-1-positive population (infected) and YOYO-1-negative population (uninfected) using a BD Accuri C6 automated flow cytometer.

### *Toxoplasma gondii* parasite culture

Parasites were grown in human foreskin fibroblasts (HFFs) using a mixture of Dulbecco’s modified Eagle’s medium (DMEM) with 10% FetalPlex animal serum complex (Gemini Biotech, catalog no. 100602), 2LmM l-glutamine, 100 μg/ml penicillin, and 100 μg/ml streptomycin. Parasites were cultured at 37°C in 5% CO_2_.

### FP-rho labeling competition assays

*Plasmodium* parasites (ring/troph, 13% parasitemia) were treated with either DMSO or salinipostin A for 3 h at 37°C. Cultures were centrifuged and RBCs were saponin-lysed (0.15%). Parasite pellets were washed with PBS and lysed in lysis buffer (50 mM Tris, pH 7.4, 150 mM NaCl, 0.5% NP40, 0.1% SDS) for 30 min on ice. Intact *T. gondii* parasites and HEK 293T cells were treated as previously described (Garland et al., 2018). Lysates were then clarified by centrifugation at 4°C, and the protein concentration was quantified using BCA. 50 μg lysate was incubated with 1 μM FP-rho for 1 h at 37°C. Reactions were quenched with reducing SDS sample buffer, and then the entire reaction was resolved by SDS-PAGE. A typhoon flat-bed scanner was used to scan the gel (532-nm laser, 610-nm filter, PMT700). Where necessary, the gel was then western blotted.

### Salinipostin alkyne labeling

*Plasmodium* parasites (ring/troph, 13% parasitemia) were treated with either DMSO or salinipostin A for 2 h at 37°C, followed by Sal alkyne for 2 hours at 37°C with light rocking. Pellets were prepared and lysed as described above. Click labeling was performed by adding 50 μg of membrane lysate in 44 μL of total volume with PBS. Add 1 μL of 50 mM CuSO_4_, 1 μL 50 mM TCEP, 3 μL of 1× ligand (Tris[(1-benzyl-1H-1,2,3-triazol-4-yl)methyl]amine (Sigma-Aldrich) 20% DMSO in *t*-butanol), and 1 μL of Rhodamine-azide. Vortex and let sit at room temperature for 30 minutes, then vortex and let sit another 30 minutes. Add SDS gel loading buffer. Run standard SDS-PAGE gel and scan gel fluorescence as above.

### Sample preparation and liquid chromatography–mass spectrometry (LC-MS) analysis for proteomics

*P. falciparum* W2 cultures (3 × 125 mL) were grown to 23% parasitemia at a ratio of 1:1 rings:trophozoites, and treated with either vehicle (DMSO), 1 μM Sal alk, or 1 μM Sal A, for 2 h at 37°C with slow shaking. For the 1 μM Sal A treatment, 1 μM Sal alk was added after the initial 2 h incubation and the flask was returned to the incubator for an additional 2 h. Parasites were harvested by saponin lysis. Parasite pellets were lysed in 50 mM TEOA, 4% SDS, 150 mM NaCl, pH 7.4 and quantified. Two milligrams of total protein per technical replicate (n=3) was subjected to a click reaction in 500 μL total volume by addition 20 μL CuSO_4_, 60 μL TBTA, 20 μL 5 mM biotin azide, and 20 μL TCEP and rotating at room temperature for 2.5 h. Samples were prepared for LC-MS analysis as described previously (Weerapana et al., 2007).

### Expression, purification of recombinant PfMAGLLP

A codon-optimized version of the PF3D7_1038900 gene was purchased from IDT (Integrated DNA Technologies Inc., San Diego, CA, USA). After cloning into the expression vector pET-28a, the protein contains an N-terminal hexa-His tag. Proteins were expressed in BL21-CodonPlus (DE3)-RIL *Escherichia coli* (Agilent Technologies, catalog no. 230245) at 37°C with 0.25 mM IPTG (Isopropyl β-D-1-thiogalactopyranoside) for 4 h. Recombinant proteins were purified as previously described (Child et al., 2013) with the following modifications: 0.02% NP-40 detergent and 10% glycerol were included in both the lysis and wash buffer. Protein concentrations were quantitated by bicinchoninic acid (BCA) assay.

### Structure prediction and modeling of PfMAGLLP

The homology model building comprises two steps: finding best structural template by performing sequence alignment of a target sequence with sequences of proteins, for which experimental structure is known, and then applying modeling program that builds three-dimensional model satisfying spatial constraints of the template structure. We applied FFAS (Fold & Function Assignment) program (Jaroszewski et al., 2011) for the alignment against PDB structures and Modeller software (Šali and Blundell, 1993) for building the model structures.

### Fluorogenic substrate assays

10 μL of a rPfMAGLLP solution (5 nM) in PBS containing 0.05% Triton X-100, 1 mM DTT were added to the wells of a black flat-bottom 384-well plate. Stock solutions of 4-methylumbelliferyl (4-MU)-based fluorogenic substrates were dissolved in DMSO (1◻mM) and diluted in assay buffer. 10 μL of substrates was added and fluorescence (λex◻=◻365◻nm and λem◻=◻455◻nm) was read at 37°C in 1 min intervals on a Cytation 3 imaging reader (BioTek, Winooski, VT, USA) for 60◻min. Turnover rates in the linear phase of the reaction were calculated using as RFU/sec. For inhibitor screening, the compounds were pre-incubated with enzyme for 30 min at room temperature prior to the addition of substrate.

### Monoacylglycerol lipase assay

To a 199 μL of enzyme (50 nM) in 10 mM Tris, 1 mM EDTA, pH 7.4 in an opaque flat-bottom 96-well plate were added 0.5 μL of arachidonoyl-1-thio-glycerol 25 mM (final concentration of 62.5 μM) and 1 μL of DTNB 25 mM (final concentration of 125 μM). The plate was carefully shaken for 10 sec and absorbance was read at 412 nm for 45 min at room temperature.

### FP-Rho labeling of rPfMAGLLP and human MAGL

rPfMAGLLP and human MAGL (50 nM) in PBS with 1 mM DTT was preincubated with different concentrations of compounds or DMSO for 30 min at room temperature, before FP-Rho (1 μM) was added. Samples were incubated at room temperature for 30 min, boiled in SDS loading buffer and analyzed by SDS-PAGE.

### Generation of overexpressing parasites (pLN and pRL2)

Pf_0818600, Pf_1038900, or Pf_0805000 were synthesized as GeneBlocks (Basler et al.) and cloned in pLN and pRL2 plasmids containing an HA tag and transfected as previously described (Gisselberg et al., 2013) using the Bxb1 mycobacteriophage integrase system in Dd2 strain parasites containing the attB recombination site in combination with a red blood cell (RBC) preloading technique. Successful integration was confirmed by PCR and Western blot.

### Imaging of PfATP4-GFP parasites

Sorbitol-synchronized parasites expressing the parasite plasma membrane protein PfATP4 fused to GFP(Rottmann et al., 2010), were treated at 24~30 hours post-invasion (hpi) with either 2 μM Sal A, 6 μM Orlistat or DMSO (control), and cultured for an additional 18 h. Cells were then harvested and fixed in 4% v/v formaldehyde supplemented with 0.0075% v/v glutaraldehyde (Sigma) in 1×PBS for 30 min at room temperature. Cell membranes were permeabilized in 0.1% Triton X-100 in PBS for 30 min and autofluorescence was quenched using 0.1 M glycine in PBS for 15 min. Blocking was performed with 3% w/v bovine serum albumin (BSA) for 1 h at room temperature. Cells were incubated with anti-GFP (Clonetech; 1:200 dilution in 3% BSA and 0.1% Tween in PBS) for 90 min at room temperature, and for 60 min with an anti-rabit Alexa Fluor^TM^ 488-conjugated secondary antibody (ThermoFisher, 1:2000 dilution in 3% BSA and 0.1% Tween in PBS). Thin blood smears of stained RBCs were prepared on microscope slides and mounted with cover slips using Prolong Diamond Antifade Mountant with DAPI (ThermoFisher). Slides were imaged using a Nikon Eclipse Ti-E wide-field microscope equipped with an sCMOS camera (Andor) and a Plan Apochromate oil immersion objective with 100× magnification (1.4 numerical aperture). A minimum of 20 Z-stacks (0.2 μm step size) were taken of each parasitized RBC. NIS-Elements imaging software (Version 5.02, Nikon) was used to deconvolve images and perform 3D-volume reconstructions. Deconvolution was performed using 25 iterations of the Richardson-Lucy algorithm for each image.

### Sample preparation for lipidomics

*P. falciparum* W2 cultures were maintained in human RBCs in RPMI 1640 complete medium and synchronized with 5% D-sorbitol (w/v) for two or more successive growth cycles and expanded to 8-10% parasitemia prior to harvesting. For lipid analysis, 75 ml cultures were prepared at 8-10% parasitemia and 4-6% hematocrit. Cultures were treated with or without 6.4 μM JW651 (8XEC_50_) at 8 hpi (triplicate cultures of one bioreplicate). Parasites were monitored by blood smears and harvested at 48 hpi while drugs were maintained until the final time point. The medium was removed and cells were resuspended in 1.5 ml of PBS buffer and transferred into glass vial pre-loaded with 2:1 CHCl_3_:MeOH to final of 2:1:1 CHCl_3_:MeOH:PBS. Samples were shaken and centrifuged at 1000 RPM for 5 min at 4◻°C. The bottom layer was pulled into a new glass vial and dried under nitrogen gas flow. Samples were then analyzed on QTOF.

### Analysis of lipids using high-performance liquid chromatography-mass spectrometry

Mass spectrometry analysis was performed with an electrospray ionization (ESI) source on an Agilent 6545 Q-TOF LC/MS in positive and negative ionization modes. For Q-TOF acquisition parameters, the mass range was set from 100 to 1200 m/z with an acquisition rate of 10 spectra/second and time of 100 ms/spectrum. For Dual AJS ESI source parameters, the drying gas temperature was set to 250°C with a flow rate of 12 l/min, and the nebulizer pressure was 20 psi. The sheath gas temperature was set to 300°C with a flow rate of 12 l/min. The capillary voltage was set to 3500 V and the fragmentor voltage was set to 100 V. For separation of nonpolar metabolites, reversed-phase chromatography was performed with a Luna 5 μm C5 100 Å LC column (Phenomenex cat # 00B-4043-E0). Samples were injected at 20 μl each. Mobile phases for positive ionization mode acquisition were as follows: Buffer A, 95:5 water/methanol with 0.1% formic acid; Buffer B, 60:35:5 isopropanol/methanol/water with 0.1% formic acid. Mobile phases for negative ionization mode acquisition were as follows: Buffer A, 95:5 water/methanol with 0.1% ammonium hydroxide; Buffer B, 60:35:5 isopropanol/methanol/water with 0.1% ammonium hydroxide. All solvents were HPLC-grade. The flow rate for each run started with 0.5 minutes 95% A / 5% B at 0.6 ml/min, followed by a gradient starting at 95% A / 5% B changing linearly to 5% A / 95% B at 0.6 ml/min over the course of 19.5 minutes, followed by a hold at 5% A / 95% B at 0.6 ml/min for 8 minutes and a final 2 minutes at 95% A / 5% B at 0.6 ml/min. The raw Agilent .d files were converted to mzXML format using MSConvert and then imported into XCMS software to identify features specifically differentially regulated in drug-treated versus vehicle-treated samples. The following XCMS parameters were used: step = 0.05, mzdiff = 0.05, snthresh = 5, mzwid = 0.1.

### *In vitro* evolution of drug-resistant lines

To select for Sal A-resistant parasites, duplicate flasks of 1×10^9^ Dd2_Polδ parasites, each with a starting parasitemia below 2%, were exposed to Sal alk at a concentration of 2.5μM (~2× the IC_90_ concentration). Drug-containing media was refreshed every other day for the first ten days, after which drug pressure was removed and cultures were continued in drug-free medium. Recrudescence was monitored microscopically.

### DNA library preparation and whole genome sequencing

200 ng of genomic DNA was used to prepare DNA sequencing libraries using the Illumina Nextera DNA Flex library prep kit with dual indices. The samples were multiplexed and sequenced on an Illumina MiSeq to obtain 300 base-pair paired-end reads at an average of 50× depth of coverage. Sequence reads were aligned to the *P. falciparum* 3D7 genome (PlasmoDB version 36) using Burrow-Wheeler Alignment (BWA). PCR duplicates and unmapped reads were filtered out using Samtools and Picard. Reads were realigned around indels using GATK RealignerTargetCreator and base quality scores were recalibrated using GATK BaseRecalibrator. GATK HaplotypeCaller (version 3.8) was used to identify all SNPs in the resistant clones. SNPs were filtered based on quality scores (variant quality as function of depth QD > 1.5, mapping quality > 40, min base quality score > 18) and read depth (depth of read > 5) to obtain high quality SNPs. These SNPs were annotated using snpEFF. Comparative SNP analyses between the resistant clones and the parental Dd2_Polδ line were performed to generate a final list of SNPs that were present exclusively in the resistant clones. Integrated genome viewer (IGV) was used to visually verify the presence of these SNPs in the clones. BIC-Seq(Xi et al., 2011) was used to check for copy number variations (CNVs) in the resistant mutants using the Bayesian statistical model. CNVs in highly variable surface antigens and multi-gene families were removed as these are prone to copy number changes with *in vitro* culture.

### *In vitro* determination of IC_50_ levels for Sal A-selected clones

IC_50_ values were determined by testing parasites against two-fold serial dilutions of antimalarial compounds (Ekland et al., 2011). Compounds were tested in duplicate in 96-well plates, with 200 μL final volumes per well. Parasites were seeded at 0.2% parasitemia and 1% hematocrit. Serial dilutions and were performed on a Freedom Evo 100 liquid-handling instrument (Tecan). Parasites were continuously exposed to drugs for 72 h. After 72 h, cultures were cut 1:5 and continued in drug-free medium for an additional 48 h to ensure complete parasite killing. After 120 h, parasites were stained with 1×SYBR Green and 100 nM MitoTracker Deep Red (ThermoFisher) and parasitemias were measured on an Intellicyt iQue3 flow cytometer sampling 10,0000–20,000 events per well. Data were analyzed using FlowJo and IC_50_ values were derived using nonlinear regression analysis (GraphPad Prism).

### Chemical synthesis of salinipostin alkyne

#### 15-hexadecyn-1-ol

1 g of NaH (42 mmol, 10 equiv.) was added to a dry round bottom flask under argon. 1,3-diaminopropane (15 mL, 180 mmol) was slowly added to the flask while stirring. The mixture was heated to 70° C and stirred for 1 h. After cooling to room temperature, 1 g of 7-hexadecyn-1-ol (4.2 mmol, dissolved in 4 mL 1,3-diaminopropane) was added to the flask and the reaction proceeded overnight at 55°C. The reaction was cooled to room temperature and ice cold H_2_O was added. The reaction mixture was acidified by the addition of 12% HCl. The organic layer was extracted 3x with ether, washed with saturated sodium bicarbonate and brine, and dried with MgSO_4_. The organic layer was dried in vacuo and the resulting off-white powder was purified by silica column chromatography using EtOAc/hexanes step gradients (0%, 5%, 10%, 15%) to afford 15-hexadecyn-1-ol as a white solid in 57% yield. ^1^H NMR (400 MHz, Chloroform-d) δ 3.63 (t, *J* = 6.6 Hz, 2H), 2.17 (td, *J* = 7.1, 2.6 Hz, 2H), 1.93 (t, *J* = 2.6 Hz, 1H), 1.59 – 1.46 (m, 4H), 1.44 – 1.23 (m, 20H). ^13^C NMR (100 MHz, Chloroform-d) δ 84.96, 68.16, 63.20, 32.93, 29.80-29.70, 29.64, 29.57, 29.25, 28.90, 28.62, 25.87, 18.53.

#### 16-bromohexadec-1-yne

To a solution of 15-hexadecyn-1-ol (150 mg, 0.63 mmol) in dichloromethane (3.7 mL) was added CBr4 (313 mg, 0.95 mmol, 1.5 equiv.) and PPh3 (248 mg, 0.95 mmol, 1.5 equiv.) and the reaction mixture was refluxed until the starting material was completely consumed by TLC analysis. The crude product was adsorbed onto celite, and purified using silica column chromatography (hexanes) to afford 16-bromohexadec-1-yne in 47% yield. ^1^H NMR (400 MHz, Chloroform-d) δ 3.39 (t, *J* = 6.9 Hz, 2H), 2.16 (td, *J* = 7.1, 2.7 Hz, 2H), 1.92 (t, *J* = 2.7 Hz, 1H), 1.83 (m, 2H), 1.57 (m, 2H), 1.44-1.34 (m, 4H), 1.31-1.25 (m, 16H). ^13^C NMR (100 MHz, Chloroform-d) δ 84.86, 68.15, 34.09, 32.97, 29.74-29.70, 29.66, 29.62, 29.56, 29.24, 28.90, 28.89, 28.63, 28.31, 18.52.

#### Salinipostin alkyne (1)

A mixture of salinipostins A, B, and C (1.2 mg, ~2.7 μmol) and 16-bromohexadec-1-yne (18 mg, 60 μmol, 22 equiv.) were solubilized in 0.4 mL anhydrous dioxane. A catalytic amount of TBAI was added and the mixture was immersed in an oil bath preheated to 110°C and refluxed for 5 h. The reaction progress was monitored by LC/MS. After consumption of the starting material, the reaction mixture was dried and purified by semi-preparative HPLC (Agilent Zorbax 300SB-C18 column, 5 μm, 9.4 x 250 mm using a gradient of ACN:H_2_O + 0.1% TFA from 70-98% over 15 min to afford two diastereomers of salinipostin alkyne. Peak 1 (Salinipostin alkyne): ^1^H NMR (601 MHz, Chloroform-d) δ 4.43 (t, *J* = 9.0 Hz, 1H), 4.30 (dd, *J* = 18.9, 6.2 Hz, 1H), 4.23 – 4.12 (m, 2H), 4.04 (dd, *J* = 9.5, 5.8 Hz, 2H), 3.83 – 3.73 (m, 1H), 2.88 (ddt, *J* = 110.4, 14.2, 7.6 Hz, 2H), 2.18 (tdd, *J* = 7.1, 2.7, 1.0 Hz, 2H), 1.94 (td, *J* = 2.6, 1.0 Hz, 1H), 1.72 (p, *J* = 6.8 Hz, 2H), 1.64 – 1.47 (m, 4H), 1.43 – 1.34 (m, 6H), 1.26 (d, *J* = 7.7 Hz, 16H), 0.93 (td, *J* = 7.3, 1.0 Hz, 3H). ^31^P NMR-δ 12.24 (s).

Peak 2 (epi-Salinipostin alkyne): ^1^H NMR (601 MHz, Chloroform-d) δ 4.49 – 4.41 (m, 1H), 4.37 – 4.29 (m, 1H), 4.24 (dt, *J* = 8.0, 6.6 Hz, 2H), 4.14 (ddd, *J* = 25.7, 11.2, 3.6 Hz, 1H), 3.83 – 3.74 (m, 2H), 2.98 – 2.86 (m, 2H), 2.18 (tt, *J* = 5.4, 2.6 Hz, 2H), 1.94 (t, *J* = 2.6 Hz, 1H), 1.74 (p, *J* = 6.7 Hz, 2H), 1.53 (s, 4H), 1.40 (dt, *J* = 15.1, 7.4 Hz, 7H), 1.27 (d, *J* = 7.3 Hz, 15H), 0.96 – 0.90 (m, 3H). 31P NMR-δ 8.88 (s).

## Supporting information

Supplemental Methods, Figures and Tablespplemental Methods, Figures and Tablesl

## ACKNOWLEDGMENTS

We thank Daniel Goldberg for sharing the parasites expressing wild-type (WT) and active site mutant (S179T) PfPARE-GFP fusions. We thank Benjamin Cravatt for providing human MAGL inhibitors and a library of triazole urea containing compounds. We also thank Ellen Yeh for access to the BD Accuri flow cytometer and helpful discussion. We thank Edgar Deu for his efforts to generate conditional knockout of PfMAGLLP. We thank Micah Niphakis and Kenneth Lum for the sequence and phylogenetic analysis of serine hydrolases and advice. Funding for this work was provided by R33 AI127851 (to M.B. and D.A.F.), R01GM117004 and R01GM118431 (to E.W.), DK105203 and P30DK116074 (to J.Z.L.), and NSERC Discovery (to R.G.L.).

## AUTHOR CONTRIBUTIONS

Conceptualization, E.Y., C.J.S., E.W., and M.B.; Methodology, J.Z.L., A.U., E.W., D.A.F., and M.B.; Formal Analysis, E.Y., C.J.S., B.H.S., T.Y., Y.Z., K.K., P.C., J.Z.L, E.W., D.A.F., and M.B.; Investigation, E.Y., C.J.S., B.H.S., O.O., T.Y., S.M., N.F.G., K.K., I.T.F., S.M.T., M.J.B., P.C., and E.W.; Writing – Original Draft, E.Y., C.J.S., and, M.B.; Writing – Review & Editing, E.Y., C.J.S., B.H.S., D.A.F., and M.B.; Resources, R.G.L., J.Z.L., A.U., E.W., D.A.F., and M.B.; Supervision, E.W., D.A.F., and M.B.; Funding Acquisition, R.G.L., J.Z.L., E.W., D.A.F., and M.B.

## DECLARATION OF INTERESTS

The authors declare no competing interests.

