## Supplemental Methods, Figures and Tablespplemental Methods, Figures and Tablesl for "The antimalarial natural product salinipostin A identifies essential α/β serine hydrolases involved in lipid metabolism in *P. falciparum* parasites"

**Table of contents**

Figure S1. Growth inhibition of *Pf* parasites by salinipostin alkyne.

Figure S2. Assessment of *Pf* parasites expressing wild-type (WT-GFP) and active site mutant (S179T-GFP) PfPARE.

Figure S3. Sequence and structure alignment of PfPARE or PfXL2 with human MAGL.

Figure S4. Structure prediction of PfMAGLLP active site and processing of 4-methylumbelliferyl (4-MU)-based fluorogenic substrates by PfMAGLLP.

Figure S5. Inhibition of PfMAGLLP by serine-reactive small molecules.

Figure S6. Effect of Sal A on the viability of *P. falciparum* parasites overexpressing HA-tagged Pf_0818600, Pf_1038900, or Pf_0805000.

Figure S7. Monoacylglycerols identified in JW651-treated parasites.

Table S1. Proteomics raw data.

Table S2. IC_50_ and IC_90_ values of *P. falciparum* lines selected for resistance to Sal A.

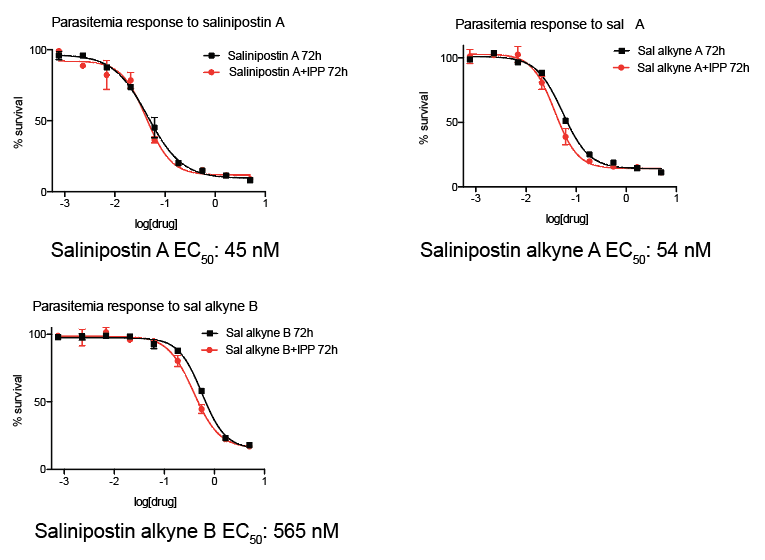

**Figure S1.** **Effect of isopentenyl pyrophosphate on inhibition of cell growth by Sal A and Sal alk. Related to Figure 2.** Growth curves of *P. falciparum* parasites in the presence of salinipostin A or salinipostin alkyne diastereomers A or B in the absence and presence of isopentenyl pyrophosphate.

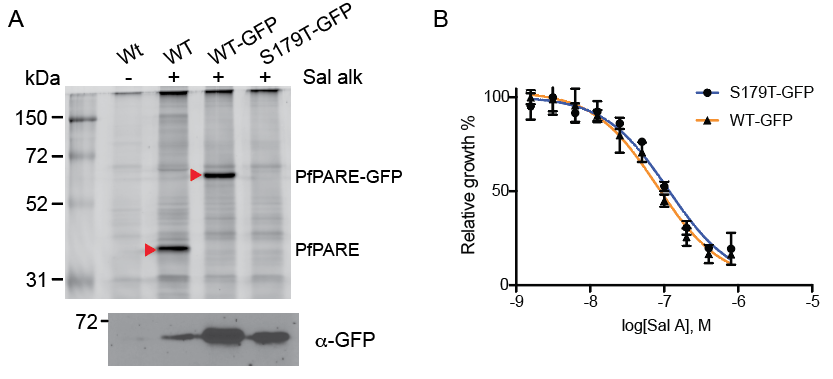

**Figure S2. Sal alk labels the serine hydrolase PfPARE. Related to Figure 2.** (A) In-gel fluorescence of *P. falciparum* parasites expressing wild-type (WT-GFP) and active site mutant (S179T-GFP) PfPARE-GFP fusions labeled by salinipostin alkyne probe. Note the shift of molecular weight for a GFP fusion of PfPARE (red arrow). (B) Potency of salinipostin A against *P. falciparum* parasites in the presence or absence of active PfPARE.

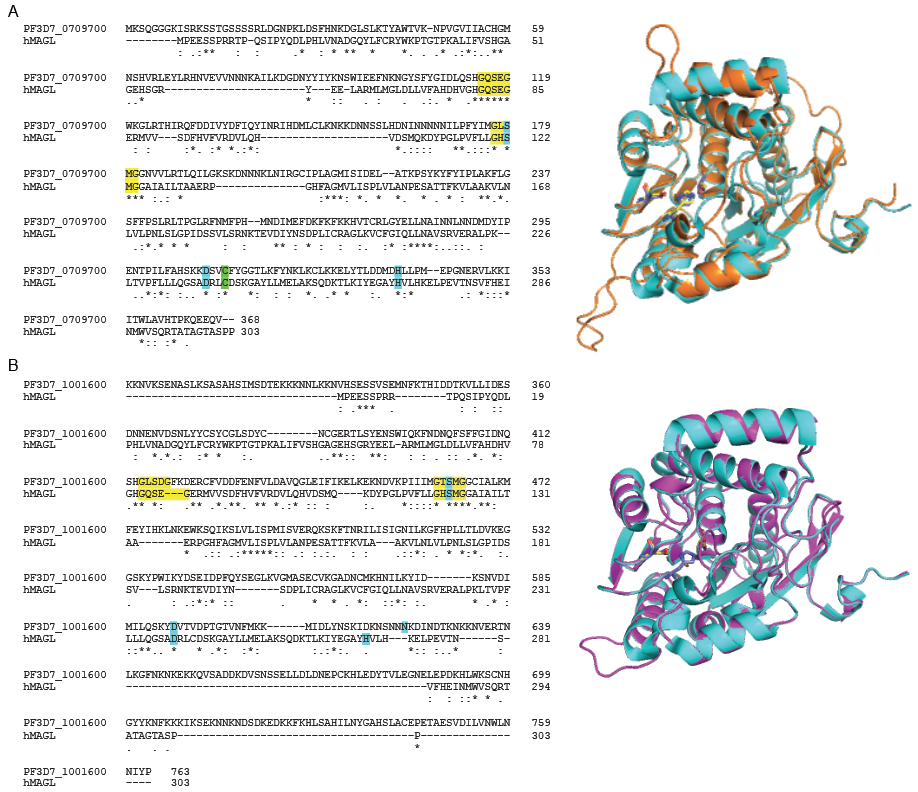

**Figure S3. Sequence and structural alignment of PfPARE and PfXL2 hydrolases with human MAGL. Related to Figure 3.** (A) Sequence of PfPARE is aligned with hMAGL using Clustal W2. Identical and highly similar residues are indicated with (*) or (:), respectively. Sequence identity is 20% and FFAS (fold and function assignment system) score is -75.4. The conserved G-X-S-X-G motif and residues of the catalytic triad are highlighted in yellow and cyan, respectively. Predicted structure of PfPARE (orange) is superimposed with human MAGL structure (PDB: 3jw8, cyan). Catalytic triads are shown in yellow for PfPARE and purple for hMAGL. (B) Sequence of PfXL2 is aligned with hMAGL. Sequence identity is 18% and FFAS score is -71.4. Predicted structure of PfXL2 (magenta) is superimposed with human MAGL structure (PDB: 3jw8, cyan).

A

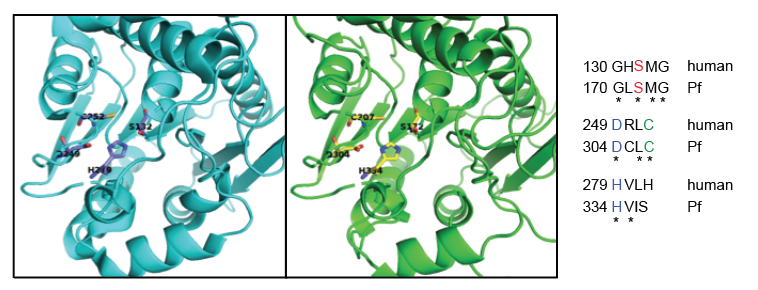

B

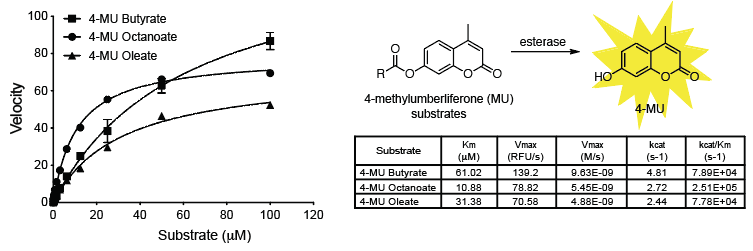

**Figure S4. The serine hydrolase PF3D7_1038900 (PfMAGLLP) aligns with human MAGL and can process lipid ester substrates. Related to Figure 3**. (A) Predicted structure of PfMAGLLP (green) based on the template structure of hMAGL (PDB: 3jw8, cyan). Active site residues are Ser172, Asp304, His334 in PfMAGLLP (yellow) and Ser132, Asp249, His279 in 3jw8 (purple). Identical residues are indicated with (*). The conserved residues of the catalytic triad are highlighted in red for serine and blue for aspartate and histidine. The cysteine adjacent to the active site in each protein is highlighted in green (relates to Figure 3B). (B) Processing of 4-methylumbelliferyl (4-MU)-based fluorogenic substrates by rPfMAGLLP. The enzyme (2 nM) was incubated with different concentrations of each substrate. Velocities for each substrate are depicted as relative fluorescence units/sec (λ_ex_ = 365 nm, λ_em_ = 455 nm). Values are the means of triplicates ± standard error. The kinetic parameters K_m_ and V_max_ were determined via nonlinear regression analysis using GraphPad Prism.

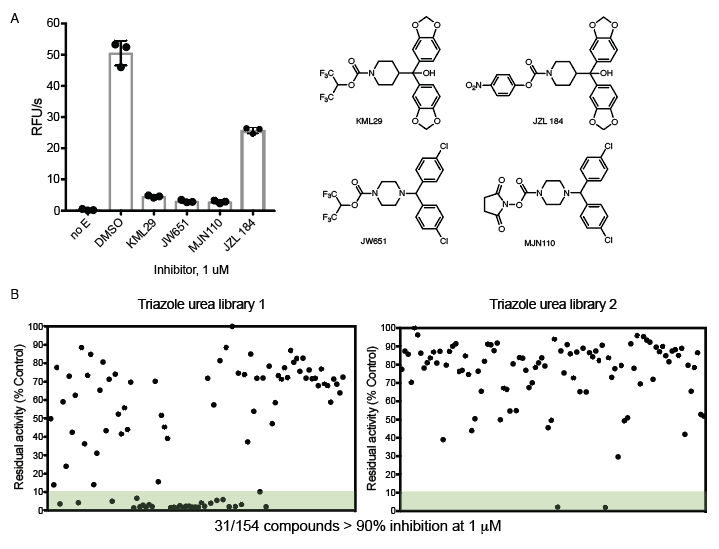

C

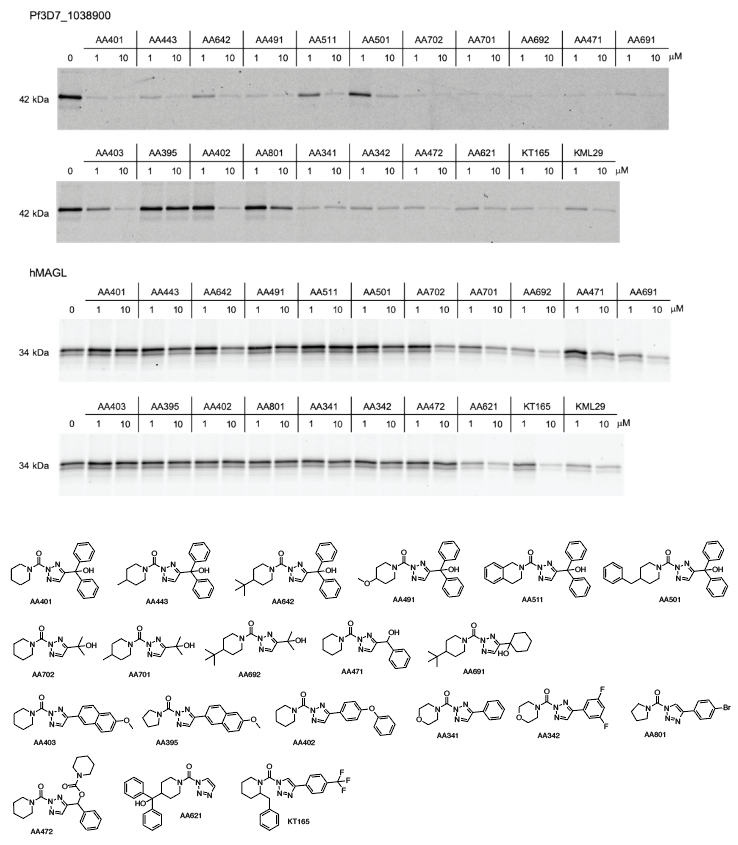

D

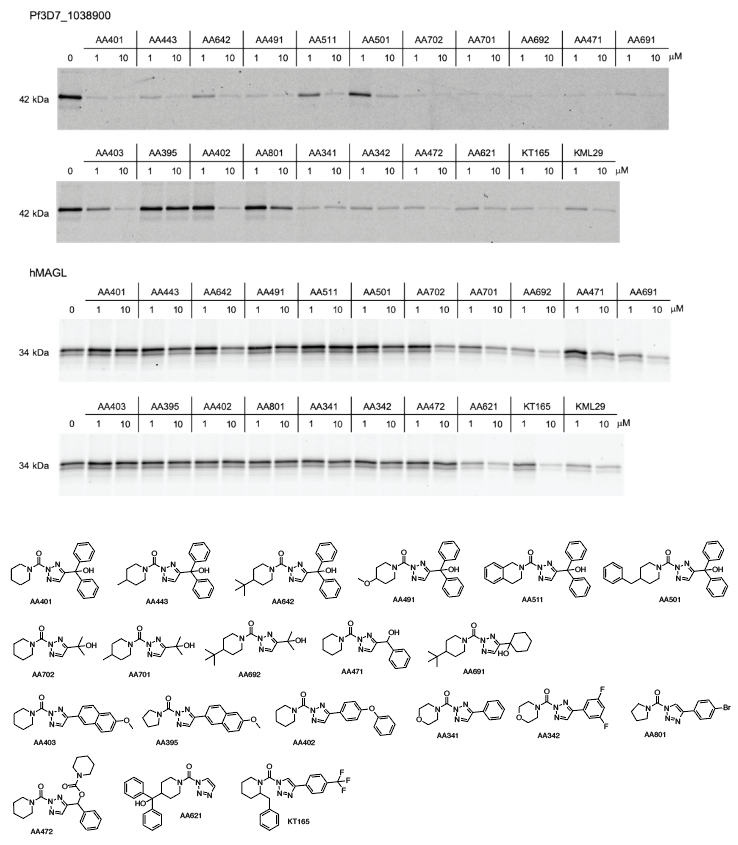

**Figure S5. Human MAGL inhibitors and triazole urea compounds identified through screening inhibit PfMAGLLP. Related to Figure 4.** (A) Inhibition of PfMAGLLP as determined by initial cleavage rates (relative fluorescence units/sec) with known hMAGL inhibitors. Reactions contained 5 nM PfMAGLLP, 1 μM inhibitor, and 10 μM substrate 4-methylumbelliferyl octanoate (mean ± 3s.d., n = 3, no E; no enzyme added). Structures of these hMAGL inhibitors are shown (relates to Figure 4A). (B) Inhibition of PfMAGLLP with a triazole urea-containing library as determined by initial cleavage rates normalized to DMSO control. Total 154 compounds were tested at 1 μM concentration. 31 compounds showed >90% inhibition. 20 selected compounds were further screened at 100 nM concentration. (C) Competitive FP-Rho labeling of PfMAGLLP and human MAGL with selected triazole urea containing compounds. (D) Structures of selected triazole urea containing compounds.

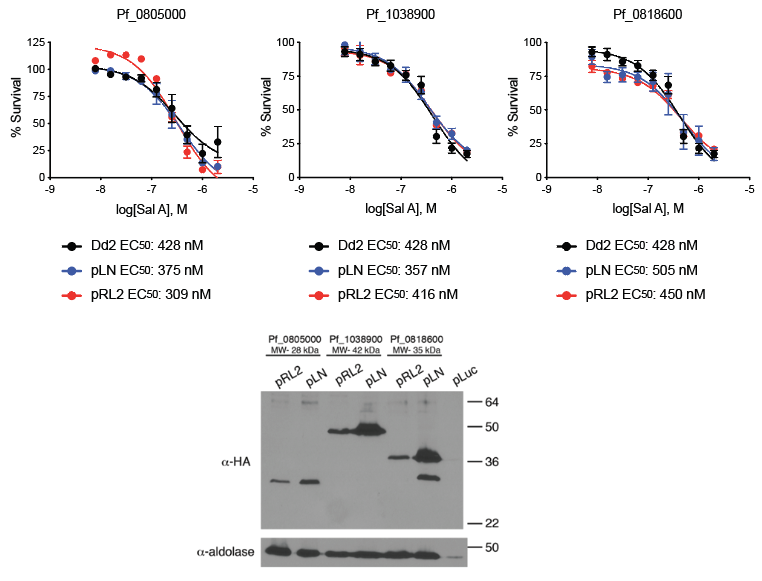

**Figure S6. Overexpression of serine hydrolase targets of Sal A do not alter dose response of Sal A parasite killing. Related to Figure 2.** Top, Effect of Sal A on the viability of *P. falciparum* parasites overexpressing HA-tagged Pf_0818600, Pf_1038900, or Pf_0805000 (pRL2; moderate, pLN; high overexpression) compared to wild-type Dd2 attB (parental line); Bottom, Western blot of expression levels of indicated proteins from the two plasmids.

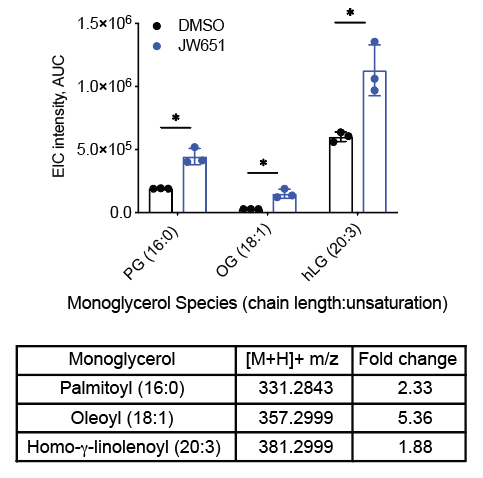

**Figure S7. The inhibitor of PfMAGLLP causes accumulation of monoacylglycerols in parasites. Related to Figure 4.** Monoacylglycerols identified in JW651-treated parasites by LC/MS analysis. Three main lipid species identified higher in JW651 treated parasites (blue) compared to DMSO treated parasites (black) were presented with chain length and number of unsaturation. Fold change was calculated based on the area under the curve (Auclair et al.) of extracted ion chromatogram (EIC).
